# A geospatial mapping pipeline for ecologists

**DOI:** 10.1101/2021.07.07.451145

**Authors:** Johan van den Hoogen, Niamh Robmann, Devin Routh, Thomas Lauber, Nina van Tiel, Olga Danylo, Thomas W. Crowther

## Abstract

Geospatial modelling can give fundamental insights in the biogeography of life, providing key information about the living world in current and future climate scenarios. Emerging statistical and machine learning approaches can help us to generate new levels of predictive accuracy in exploring the spatial patterns in ecological and biophysical processes. Although these statistical models cannot necessarily represent the essential mechanistic insights that are needed to understand global biogeochemical processes under ever-changing environmental conditions, they can provide unparalleled predictive insights that can be useful for exploring the variation in biophysical processes across space. As such, these emerging tools can be a valuable approach to complement existing mechanistic approaches as we aim to understand the biogeography of Earth’s ecosystems. Here, we present a comprehensive methodology that efficiently handles large datasets to produce global predictions. This mapping pipeline can be used to generate quantitative, spatially explicit predictions, with a particular emphasis on spatially-explicit insights into the evaluation of model uncertainties and inaccuracies.

## Introduction

Geospatial modelling is emerging as an indispensable tool in ecology. It can provide fundamental predictive insights into the environmental drivers of biodiversity, biomass and abundance that collectively determine the biogeography of life on Earth. As such, the output from such geospatial models have countless applications, including the identification of biodiversity hotspots for nature conservation [1], to gain insight in the ecological niches and tolerances of species [2], or to inform carbon or nutrient cycle models to test climate feedbacks [3]. With the advent of machine learning, larger and larger datasets can be processed, revealing previously unidentified relationships and biogeographic patterns. Although these statistical models cannot represent the critical mechanistic insights that are captured in process-based models, they can capture and describe ecological patterns with remarkable accuracy. The unparalleled predictive capacity of these emerging tools makes these geospatial models a useful complement to existing mechanistic approaches as we aim to understand the biogeography of Earth’s ecosystems.

Despite the potential for these models to provide fundamental biogeographic insights, as with any model product, a critical assessment on the prediction accuracy and model uncertainties is essential to the appropriate interpretation of results. This is particularly true for machine learning models, given their limited capacity of extrapolation and tendency to overfit [4]. At present, many large-scale geospatial modelling studies have an overall focus on single-value accuracy metrics such as root-mean-square error (RMSE), mean absolute error (MAE) or the coefficient of determination R^2^, which give a single overall measure of model uncertainty. However, such metrics only inform about the overall accuracy of the model but fail to provide quantitative spatially explicit insights into the model inaccuracies that are vitally important for assessing confidence of predictions in different regions. In addition, these accuracy measurements cannot identify the specific source of those uncertainties, which can be interactively driven, and obscured, by a wide range of spatial and statistical issues, including spatial autocorrelation, co-linearity and overfitting. It is critically important to know *where* and *how much* a spatial model is extrapolating vs interpolating, and that spatial prediction maps are only truly informative if they contain detailed spatial information about their uncertainty.

However, as with any model product, a critical assessment on the accuracy and uncertainties is essential. This is particularly true for machine learning models, given their limited capacity of extrapolation and tendency to overfit [ 4]. Although many current large-scale mapping studies have an overall focus on single-value accuracy metrics, e.g., these metrics only inform about the overall accuracy of the model but fail to provide spatial insight into the model inaccuracies. In addition, these overall accuracy measurements cannot identify the specific source of those uncertainties, which can be interactively driven, and obscured, by a wide range of spatial and statistical issues, including spatial autocorrelation, co-linearity and overfitting. Here, we argue that it is important to know *where* and *how much* a spatial model is extrapolating vs interpolating, and that spatial prediction maps are only truly informative if they contain detailed spatial information about their uncertainty.

Here, we present a general methodology (hereafter referred to as the Geospatial Mapping Pipeline or GMP) that can be used to create global high-resolution (i.e., default: 30 arc-second, or approximately 927 meters at the equator) predictions of ecological attributes, but also to generate a set of high resolution maps and metrics that allow users to evaluate the extent of extrapolation, bootstrapped confidence intervals, standard deviation, and derivatives in a spatially-explicit way. By automating this process, we hope that of the GMP pipeline’s output can help with exploratory data analyses, which are to be complimented with further analyses to explore the drivers of global biogeographic patterns. Along with providing an efficient, and robust methodology for exploring geospatial patterns, this methodology allows for a more transparent evaluation and reporting of model outputs and can aid in the interpretation of the results. The methodology requires only basic local computing power and runs fully automated in the cloud.

## Methods

### Getting started

The GMP utilizes runs on Google Earth Engine (GEE) [5] using the Python API. In order to use GEE, registration is required at https://signup.earthengine.google.com. The GEE syntax, available in Javascript and Python, is well-documented online: https://developers.google.com/earth-engine/guides.

The GMP requires Python 3 installed on a local machine, with the following modules: `ee, pandas, numpy, subprocess, time, datetime, scipy, sklearn, itertools, pathlib, math, matplotlib`. Recommended but not strictly required is `gsutil`, to facilitate the file upload processes. Note, however, that this requires access to Google Cloud Storage, which is a paid service. The corresponding steps in the approach can be replaced by manual file uploads. As all computational processes are performed by GEE, no high-end in-house computational infrastructure is required.

The purpose of the GMP was to create a modular mapping pipeline that requires no manual intervention, to facilitate automation and minimize manual error. After configuration, the GMP can automatically proceed through all necessary steps towards generating and evaluating the spatial predictions from geospatial mapping analyses. The default output includes a multiband image with the following layers: the classified image (the predicted map), bootstrapped maps (mean, confidence intervals, standard deviation and coefficient of variation), and two maps of interpolation *vs.* extrapolation, and model accuracy metrics and variable importances. The approaches to produce these output data are described in detail below. We use a global soil nematode abundance dataset [6] for illustration purposes. Leveraging GEEs cloud computing power, the GMP is relatively quick; although the total runtime depends on the project-specific configuration (e.g., the number of observations, the number of predictor layers in the composite image, the resolution of the final image, etc.), generally all calculations and export maps should be ready within 2 to 12 hours. By default, all files are exported as GEE assets. Per the user’s preferences, tables and images can be exported as csv and png or tif files, respectively.

### Generating a spatial prediction

The goal of the GMP is to take collections of measurements from discrete locations across the globe, explore the spatial and environmental patterns between those points, and then extend those patterns to predict those values in every corresponding location across the globe.

To create a spatial prediction from individual point collections of raw data, the GMP uses Random Forest (RF) models to leverage the relationships of ecological characteristics with environmental conditions, such as climate conditions, soil physiochemical properties, solar radiation or topographic characteristics (**Fig. 1**). Many of these environmental conditions are described in detail and available as spatial products, often referred to as raster layers. Such a raster layer is a gridded, spatially-explicit image where each pixel represents the value of the described property. Both raw measured data (e.g., remote-sensed products like MODIS or Landsat surface reflectance) and modelled products (e.g., most climate and soil physiochemical products) can be represented by raster data. The first preparatory step towards creating a spatial prediction is assembling a stack of representative raster layers as a composite image in GEE; these layers are used as predictor layers in the model. We provide an example composite image that contains bands describing climate, soil physiochemical, vegetation and topographic characteristics, and pixel coordinates and biome information [7–14]; see Data Availability.

**Figure 1.**
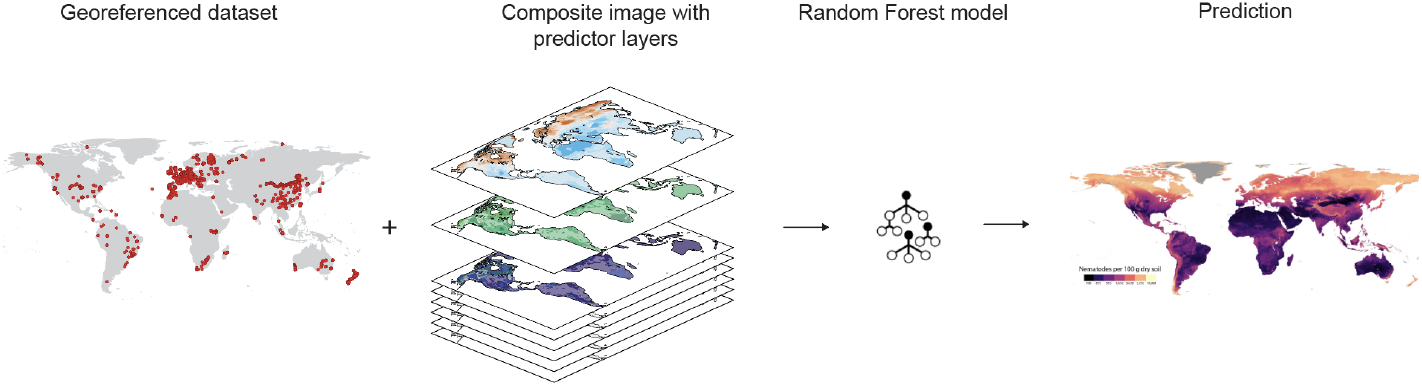
Overview of methodology. With a georeferenced observational dataset as input, the GMP uses a composite image to extract environmental information to train a RF model to create a spatial prediction.

The raw input data for the GMP are point measurements as a georeferenced dataset (i.e., a csv file with the response variable and coordinate information). Please ensure that all coordinate information is in the same coordinate reference system (default: WGS 84 EPSG:4326) and reproject if necessary. After uploading point data to GEE, pixel values for each point in the dataset are extracted from the composite image. Next, the data is aggregated at the pixel level to produce a training dataset. This is a necessary step, as duplicate pixel values can lead to overfitting and/or spatial autocorrelation (SAC) After uploading this training dataset to GEE, a set of RF models are trained and evaluated. In this process, RF hyperparameters (i.e., number of variables per split, minimum leaf population) are tuned using a grid-search procedure, and models with different combinations of hyperparameters are evaluated using random k-fold cross-validation. Cross-validation folds are assigned stratified per biome to ensure equal representation of each major bioclimatic zone in each fold. The final model, which will be used to classify the composite image to create the prediction map, is an ensemble average of the top *x* models (default: *x* = 10) can be used to create the prediction map. The benefit of this ensemble averaging approach is that it reduces the potential bias of any single prediction, albeit that this might come at the cost of a slight reduction in predicative power and increased computation time. Alternatively, the final model can be selected as the single best model from the grid search procedure based on R2, root mean square error (RMSE) or mean absolute error (MAE).

By default, the GMP assumes a global extent of the final prediction (i.e., all continents except Antarctica), at a resolution of 30 arc-seconds. Per the user’s requirements, predictions can be clipped to a continent or region of interest. Depending on the use case of the predicitons or or when the environmental predictor raster layers are available at a different resolution, the output maps can be scaled accordingly.

### Bootstrapped spatial confidence intervals

To create statistically valid per-pixel confidence intervals and uncertainty estimates, the GMP includes a stratified bootstrapping procedure (**Fig. 2**). Bootstrapping is a statistical technique that simulates sample distribution to assign measures of accuracy to model predictions (ref). In this analysis, the training dataset is sampled with replacement, using biome as stratification category. By performing a stratified sampling procedure, each biome is represented proportionally in each of the bootstrap samples. Each of these bootstrap resamples (default: *m* = 100 resamples) are then used to classify the composite image. Next, these *m* images are used to calculate mean, standard deviation and non-parametric lower 2.5% and upper 97.5% confidence interval images. We emphasize that these are the standard deviation and the 95% confidence interval of the mean value. Therefore, a particular observation may fall outside this confidence interval, but the mean of all resamples is within this confidence interval. The standard deviation and confidence interval images provide per-pixel estimates of confidence and can be used to identify regions that have markedly higher uncertainty- and hence regions where the model predictions should be taken with extra caution. These maps can also be used to calculate statistically valid upper and lower bounds of predicted values.

**Figure 2.**
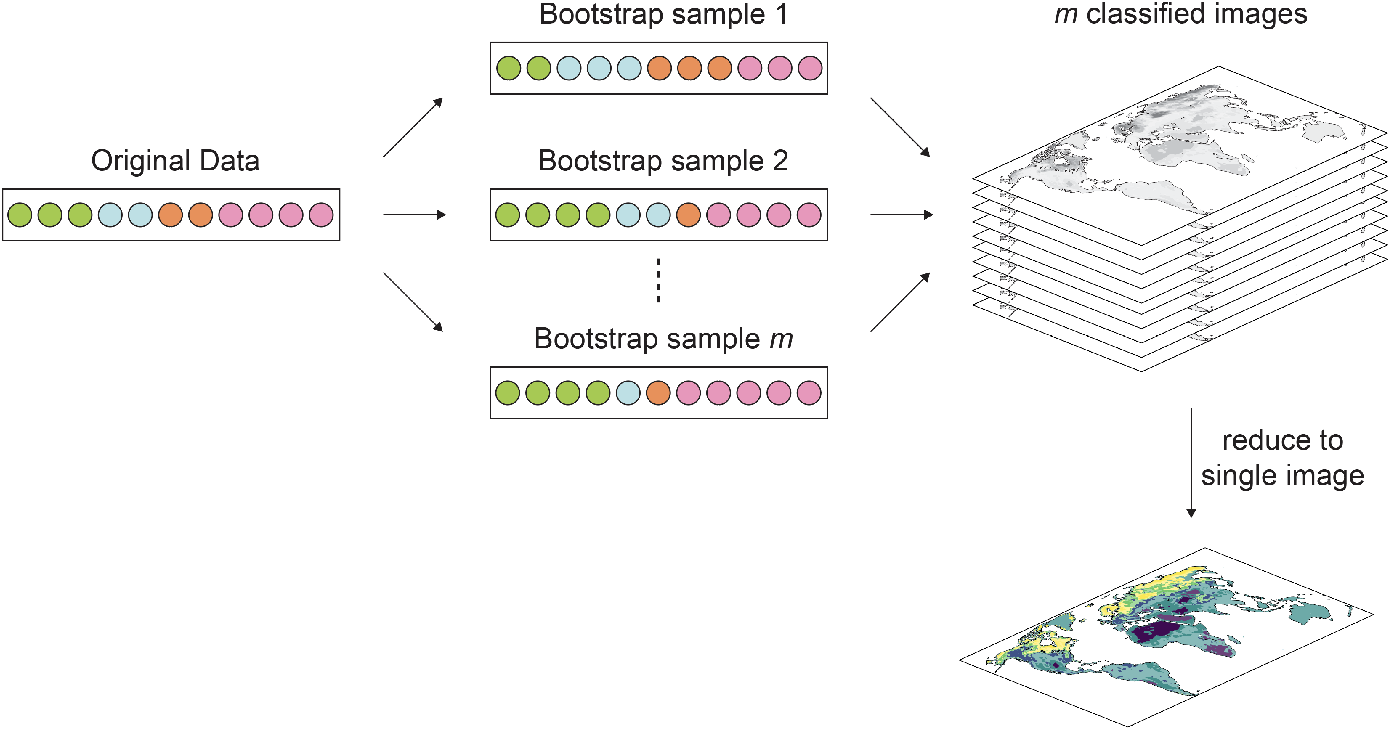
Bootstrap approach. The training dataset is sampled with replacement, with biome as stratification category. Each of these bootstrap resamples are then used to classify the composite image, and these classified images are used to calculate mean, standard deviation and lower 2.5% and upper 97.5% confidence interval images.

### Spatial assesment of extrapolation

A limitation of RF models is their limited capacity of predicting outside the range of the training data [4]. With no mechanistic understanding of the relationships between variables, extrapolating into new environments should be done with caution, as it is likely to be characterized by considerable model uncertainty. However, when making spatial predictions, especially at larger scales, it is often unavoidable that a proportion of the predictions are performed in unknown environmental space. Even with a very large dataset and a large number of environmental predictor variables, certain environmental conditions, or a specific combination of interactions between these environmental conditions, might still be outside the space represented the training data. Hence, it is critical to perform a spatial assessment of interpolation vs extrapolation, to identify regions that fall outside the training data. The GMP includes two approaches to do so: i) extrapolation in univariate space and ii) extrapolation in multivariate space.

First, to test the extent of extrapolation in univariate space we assess whether pixel values are within or outside the range the training data, this process is repeated for each environmental layer in the model (**Fig. 3**). The final image represents the proportion of bands where the pixel value falls within the range represented by the training data. A possible limitation of this method is that possible interactions between environmental predictors are not taken into account. However, assessing all possible interactions between all predictors in every pixel can result in a computationally extremely demanding task. To overcome this, we first transform the multiband composite image and training dataset into the same Principal Component (PC) space. Next, the first n PC axes are selected, collectively explaining more than *x* % variation. Then, for each bivariate combination of the selected PC axes, each pixel in the composite is tested whether it falls within the respective convex hull enclosing the training points (**Fig. 4**). The final image represents the proportion of bands where the pixel value falls within convex hull enclosing the training data points.

**Figure 3.**
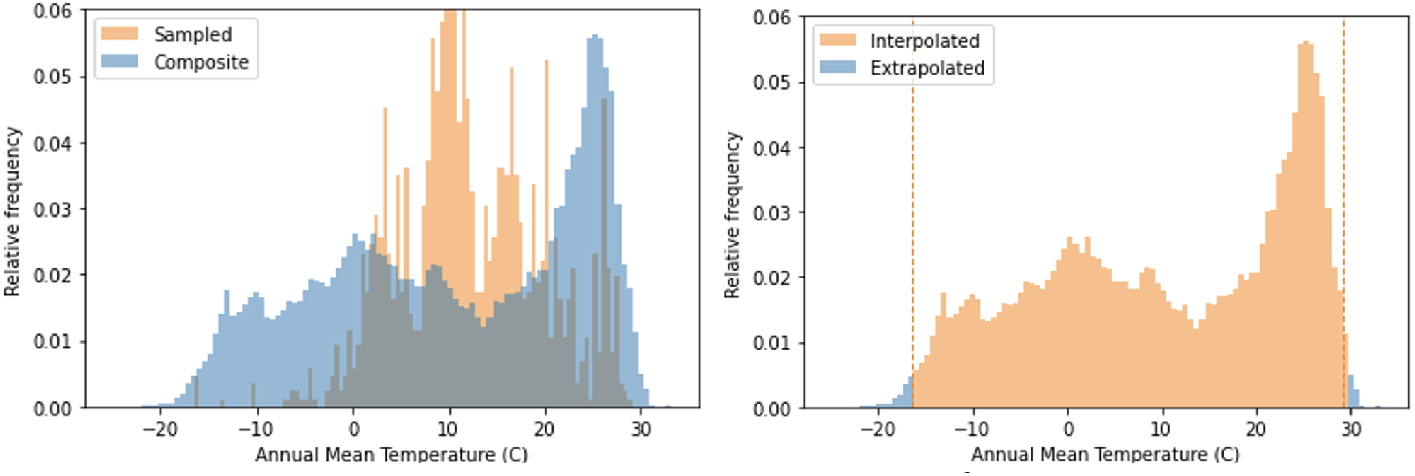
Visual representation of interpolation-extrapolation classification of composite pixels in univariate space. All pixel values of a covariate band (here: Annual Mean Temperature **(A)**) are extracted and assessed whether they fall within the minium/maximum range of the training data (dashed lines; pixel values falling within that range are classified as interpolated, those outside as extrapolated **(B)**. This process is repeated for all covariates included in the model, or alternatively, for the top *n* covariates from the variable importance analysis. The final image is the proportion of covariates that are classified as interpolated.

**Figure 4.**
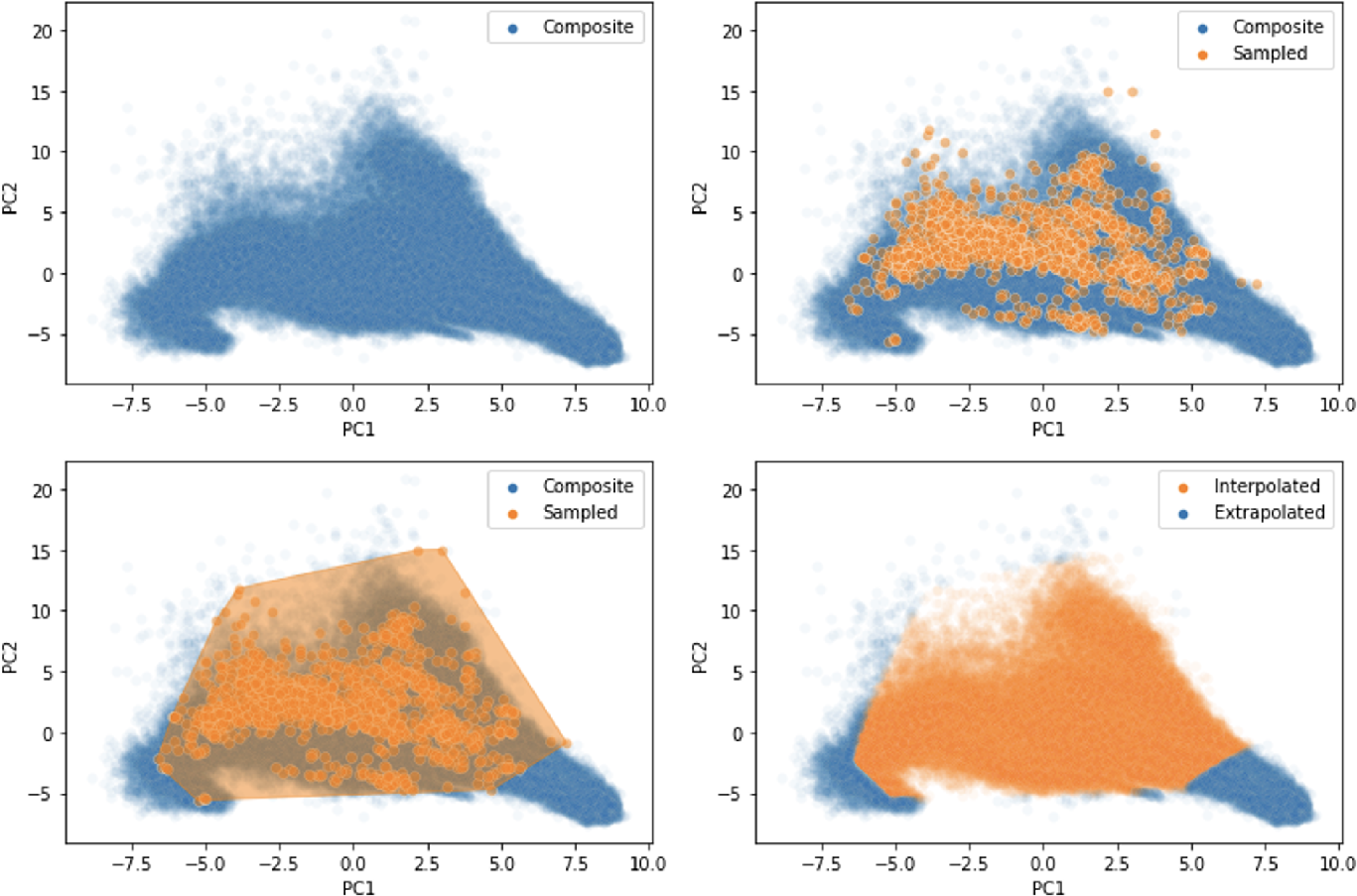
Visual representation of interpolation-extrapolation classification strategy of composite pixels in semi-multivariate space. After transforming pixel values **(A)** and sampled data **(B)** into PC space. Composite pixels falling within the convex encompassing the sampled data **(C)** are classified as interpolated, those outside as extrapolated **(D)**. This process is repeated for all bivariate combinations of the selected PC axes. The final image is the proportion of covariates that are classified as interpolated.

In our example dataset, 10 PC axes explain 90% of variation represented by the dataset. With 10 PC axes, a total of 45 bivariate combinations1 are to be taken into consideration; these combinations include PC1 × PC2, PC1 × PC3, PC1 × PC4, […], PC9 × PC10. For each of these 45 combinations, every pixel in the composite image (**Fig. 4A**) is scored to fall within or outside the convex hull enclosing the training data within that PC combination space (**Fig. 4B,C**). Pixels falling inside the convex hull are classified as interpolated (**Fig. 4D**, orange points), pixels falling outside the convex hull are classified as extrapolated (**Fig. 4D**, blue points). These classifications are summed and converted to a percentage, which is used as a proxy for the extent of interpolation vs. extrapolation. Using these maps, regions that exceed a certain extrapolation threshold can be masked in the final prediction. These maps of extrapolation can also serve as guidelines for focussed data collection efforts.

### Accounting for possible spatial autocorrelation

In many, if not most, real-world scenarios, the spatial distribution of the sampling locations is not evenly distributed. Often sampling locations are clumped together, and, as a consequence, the training data might show spatial structures that can affect the model’s performance or its accuracy assessment [15]. Specifically, when these spatial structures lead to dependence structures in the model residuals, model accuracy metrics might be overoptimistic which can lead to a false interpretation of the results.

There are a number of ways to deal with these potential biases in the dataset. First, the training data should be aggregated at the pixel level to ensure that there are no duplicate points (with equal or similar observed values), as this might lead to the model overfitting this subset of points. Next, assigning cross-validation folds stratified by bioclimatic zones (e.g., biomes, as used in the above example) can improve equal representation of environmental conditions within each fold.

To account for the potential of inflated R^2^ values due to SAC, the GMP includes a spatial leave-one-out cross-validation (SLOO-CV) test (**Fig. 5**)[15, 16]. In this test, a model is trained on all data except for points that fall within a predefined buffer zone radius *d* from the test point. This process is repeated for 1,000 randomly selected points, across each of the ten models in the final ensemble. The buffer zone radius *d* can be selected using a SAC test like Moran’s I [17] or using a semivariogram [16]. Alternatively, the spatial-leave-one process can be performed across a range of buffer zone radii.

**Figure 5.**
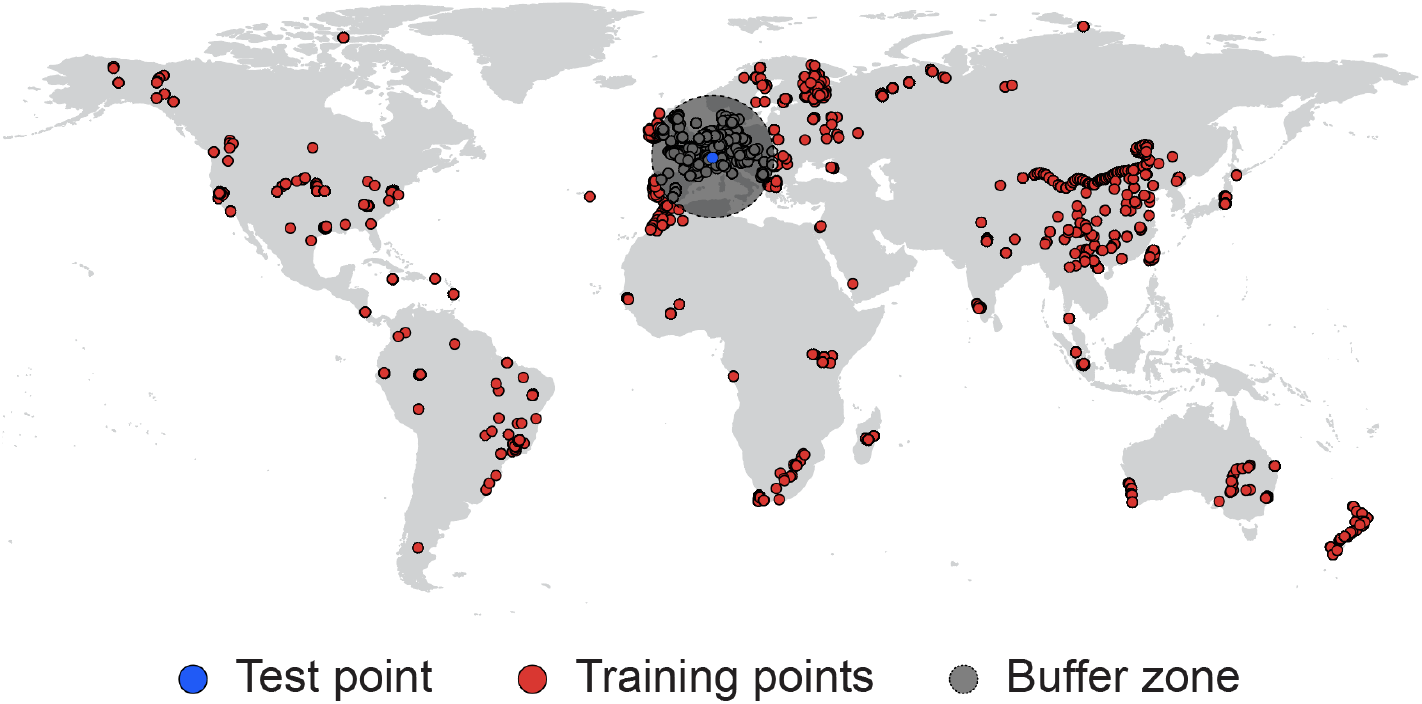
Spatial leave-one-out cross-validation approach. One point (blue) is used for testing, points that fall outside a predefined buffer zone (grey) from the test point are used for training ( red). This procedure is repeated for 1,000 randomly selected testing points, across each of the ten models in the final ensemble.

Optionally, the SLOO-CV test can be configurated as such that test points that fall outside the environmental space covered by the training data (after removing points in the buffer zone) are excluded from the accuracy assessment, thereby reducing extrapolation, in line with the methodology of [16]. However, as a result, only points that are close to one another in environmental space are taken into account. In many real-world datasets, particularly global datasets, this can result in omission of point locations or even entire regions that represent unique environmental conditions and hence these points or regions end up never being tested. As a consequence, the model performance assessment might in fact still be overoptimistic. By including points that fall outside the environmental space covered by the training data, a more conservative estimation of the model accuracy is provided.

### Discussion & limitations

The GMP provides a standardized, automated approach to generate spatially explicit maps and uncertainty estimates from point data collected across broad spatial scales. Given that geospatial mapping currently requires a number of discrete steps, it can be prone to methodological challenges that can drastically impact final outputs. By standardizing the approach, the GMP minimizes the chances of human error in generating standardized models, extrapolations and uncertainty estimates.

Yet, as with any methodology, there are a number of limitations to the GMP. First and foremost, the modelling approach is based on correlations of the response variable with environmental conditions. While Machine Learning algorithms such as RF are often able to detect previously unidentified nonlinear relationships and can provide unparalleled predictive power that is valuable to generate a fundamental understanding of Earth’s ecosystems, they are unable to provide the mechanistic insights that are required to discern the driving mechanism. Future efforts should focus on integrating these two complementary approaches.

While large datasets are required to train the RF models to accurately describe the biological variation, there is a maximum number of observations that can be used. Although this number depends on the project-specific configuration, for example the number of covariate layers, or the number of trees in the RF, generally speaking there is an upper limit of about 25,000 observations. When modelling very large datasets a (stratified) subsampling could be advisable.

We also stress that, in this current version, the GMP does not account for all sources of data uncertainty. For example, measurement uncertainty, within-plot or within-pixel variation, or other measurement-related sources of variation can influence the accuracy of the predictions. Considerable uncertainty in geospatial analyses can come from inconsistencies in data collection. Different methodologies or sampling techniques can give rise to striking differences that are not reflective of the natural variation. In each case, great care should be taken when collecting, standardizing and cleaning raw data for use in geospatial models. Our approach can help to evaluate some of the uncertainty resulting from such limitations, but it is not able to correct for and overcome these methodological inconsistencies in generating global predictions. All raw data uploaded to the pipeline will thus be taken as the same unless stated otherwise.

Another source of uncertainty is the quality of the predictor layers. As many of predictor layers are model outputs themselves, modelling errors can be propagated into the output from the GMP. Currently, these types of errors are not taken into account. Increased reporting of uncertainty of global raster layers and ecological data could facilitate additional ensemble modelling that could ultimately allow for propagation of observation and covariate uncertainty and allow users to decompose the relative contributions of different sources of uncertainty to model predictions.

## Conclusion

Altogether, the GMP offers a comprehensive and accessible approach to create high-resolution maps at broad spatial scales. By providing per-pixel estimates of uncertainty and insights into the extent of extrapolation, the GMP goes beyond the typical focus on pure predictive power. Although useful when comparing models with one another, metrics like the coefficient of determination R^2^ fail to provide insights into the spatial variation of predictive uncertainty. Moreover, as extrapolation is one of the main pitfalls of modelling approaches based on machine learning algorithms, and extrapolation is nearly unavoidable when creating global wall-to-wall maps, it is critical to know where the model is predicting outside the training space. We believe that spatial predictions can only be properly evaluated when accompanied by spatially-explicit information.

We encourage researchers from across disciplines, whether experienced biogeographers or ecologists who want to explore their data, to build upon and expand on this framework.

## Data and code availability

The source code of the GMP and associated scripts are available online: https://github.com/hooge104/geospatial_mapping_pipeline. This repo also contains a quick rundown of the necessary steps to get started. An example composite image is available as a GEE asset https://code.earthengine.google.com/?asset=users/crowtherlab/References/example_composite_30ArcSec. Data from a global soil nematode abundance dataset [6] was used to produce the figures.

## Acknowledgements

We thank Noel Gorelick, Colin Averill and Dan Maynard for insightful discussions. This research was supported by a grant from DOB Ecology to T.W.C.

## Footnotes

1 Generic formula to calculate the number of bivariate combinations: 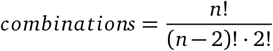

## Notes

### Competing Interest Statement

The authors have declared no competing interest.

